# BioAFMviewer AFM Analysis Arena A^3^ – an interactive software platform for automated analysis of HS-AFM imaging

**DOI:** 10.64898/2026.09.18.752612

**Authors:** Romain Amyot, Soma Yamamoto, Hiroki Konno, Clemens Franz, Holger Flechsig

## Abstract

High-speed atomic force microscopy (HS-AFM) experiments provide unique insights into nanoscale biological processes by directly visualizing the surface topography dynamics of biomolecules, such as proteins, over extended periods of time and generating large imaging data sets. However, the lack of computational methods for automated data analysis has thus far limited the full utilization of these data for the quantitative understanding of single-molecule dynamic. Here, we present the AFM Analysis Arena (A^3^) within BioAFMviewer as a software platform for automated, high-throughput quantitative analysis of AFM topographic imaging data within a user-friendly interactive graphical interface, implementing a convenient workflow from raw HS-AFM (AFM) imaging to statistical analysis. In several applications to HS-AFM imaging data, we demonstrate methods of automatized molecule detection and dynamic tracking to highlight the available capacities, aiming to transform HS-AFM measurements from observations towards an analytical tool in nanoscale biology.

## 1. Introduction

High-speed atomic force microscopy (HS-AFM) has established itself as a powerful technique in single-molecule biophysics, enabling the direct visualization of protein dynamics during functional activity [1,2]. By rapidly scanning the molecular surface with a cantilever probing tip and recording topographical changes in real time under liquid conditions, HS-AFM can observe a large number of molecules in their continuously changing functional states over extended timescales to accumulate thousands of topographic images within a single experiment. With the latest technological advancements recording 100 frames per second [3,4], the amount of available imaging data even increases by an order of magnitude.

To translate this enormous amount of valuable qualitative information into quantitative understanding of nanoscale biological processes from measurements demands the application of computational methods for automated analysis.

Indeed, most studies do not fully exploit the explanatory power of HS-AFM because high-throughput analysis methods are lacking so far. The analysis is still often performed for a confined set of data involving biased human intervention, or employing specific laboratory-developed scripts which depend on other software and remain inaccessible for broad use by the AFM community.

To overcome such shortcomings, several tools for the quantitative analysis of AFM data have been developed in the past years. Topostats [5,6] implements the computational framework for image segmentation and statistical evaluation of AFM imaging data, but it is primarily designed for the analysis of string-shaped molecules such as nucleic acids and does not provide an interface for visualizing the analysis results. ImageJ Fiji [7] is a well-established image analysis platform that provides a broad range of tools for image processing, segmentation, particle detection, tracking, and quantitative measurements. It can, in principle, be applied to HS-AFM movies but there is no direct pipeline from raw data, and it requires prior conversion to a compatible image format, such as multi-image TIFF, resulting in the loss of metadata associated with HS-AFM movies stored in their native formats. NanoLocz [8] provides an interface which can process various native AFM file format including HS-AFM formats. Like ImageJ Fiji, it provides methods for molecule detection and tracking allowing for quantitative analysis. However, NanoLocz is primarily developed for the implementation of Localization AFM [9] and currently lacks the convenient use for quantitative analysis and visualization of results.

Here, we present the AFM Analysis Arena (A^3^) within the BioAFMviewer [10] as a software platform allowing for automated high-throughput quantitative analysis of AFM topographic imaging data within a user-friendly interactive graphical interface, implementing a convenient workflow from raw HS-AFM (AFM) imaging to statistical analysis.

In several applications to HS-AFM imaging data we demonstrate methods of automatized molecule detection and dynamic tracking to highlight the available capacities, aiming to transform HS-AFM measurements from observations towards an analytical tool in nanoscale biology.

## 2. Methods and Applications

### 1. AFM Analysis Arena A^3^ software interface

The A^3^ interface consists of an AFM movie viewer, including a file manager to organize imported experimental data, an AFM data control panel for pre-processing raw data, and the analysis arena providing graphical output of analysis results. The AFM movie viewer supports import of experimental raw data in native HS-AFM data format files (.asd, .esd), as well as .jpk and multi-image TIFF files. Basic image processing, including Gaussian filtering and tilt correction, can be performed using the AFM data control panel, with the implementation previously described [11]. The analysis arena provides the interactive interface for visualization of results obtained from automated analysis and data export.

Fig. 1 shows a snapshot of the software interface highlighting the main features for automated analysis of AFM imaging data. Their applications are demonstrated in several examples as described below. For a detailed explanation of the user interface capacities, we refer to the Methods section.

**Fig. 1:**
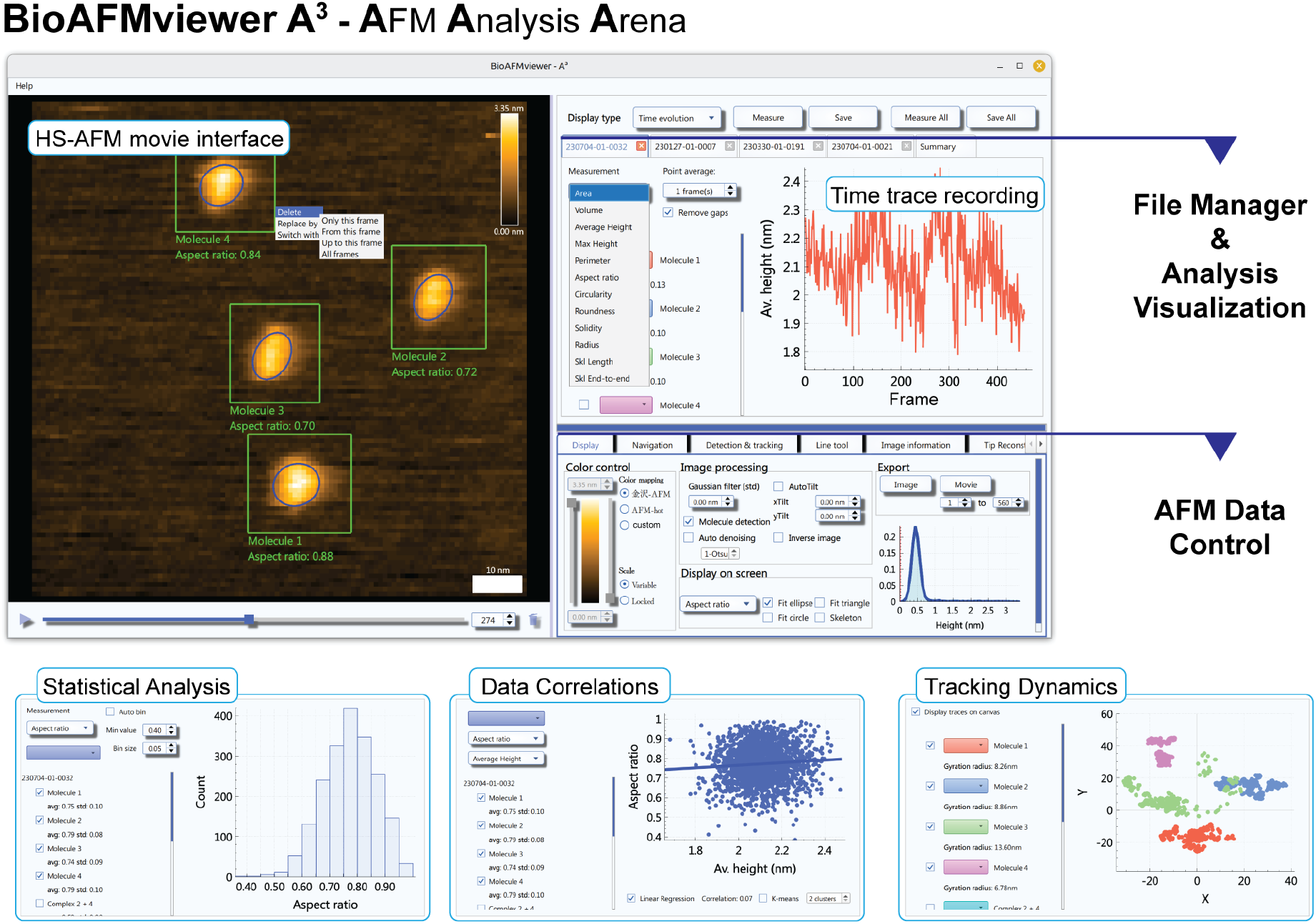
AFM Analysis Arena A^*^. Snapshot images of the software interface highlighting the major features for automated high-throughput analysis of AFM imaging data.

### 2. Automated molecule detection from HS-AFM imaging

HS-AFM topographic images must be viewed in the context of the underlying measurement procedure. They contain intrinsic imaging background coming from various AFM substrates used in experiments; the difference between most used Mica versus Aptes-coated Mica surfaces was demonstrated before [12]. On the other side, the measurements of biomolecular surface are prone to be affected by the imaging procedure itself. E.g., tip parachuting at high scanning velocities and the observation of highly dynamic samples by time-lagged imaging result in distorted topographies. Additionally, tip contamination during prolonged measurements affects imaging quality.

In HS-AFM images detection heights of biomolecular surfaces are typically at the range of several nano meters, well separated from background noise. Hence, a feasible assumption is that molecules can be isolated by the application of height thresholding to measured images. For automatized thresholding we use the Otsu method [13], which we have previously applied to HS-AFM imaging in the context of flexible fitting to identify target molecules by automation [11]. This method is also implemented in the NanoLocz software [8]. Here, we extend our previous implementation to the large-scale quantitative analysis of HS-AFM topography. As a first application we consider a data set of new experiments capturing conformational dynamic of a HECT domain protein, whose functional dynamics is involved in a cascade of processes facilitating the post-translational ubiquitination of other proteins [14]. Previous HS-AFM imaging has shown frequent switching of HECT topography shapes between oval and round shapes, indicating the occurrence of functional transitions, but the analysis of several hundreds of images to discriminate shape was done entirely by visual inspection and for a small set of selected molecules [15], hampering a solid interpretation of the acquired data.

The new data set covered the observation of 20 individual HECT domain molecules and a total of 13,432 imaging frames. For general quantitative topography analysis, we implemented geometry fitting and various shape descriptors summarized in Table T1. In the case of HECT analysis ellipse fitting and the aspect ratio was employed to evaluate topography dynamics. Fully automated molecule detection and dynamic shape quantification for a selected HECT domain data set is demonstrated in Supplementary Movie V1, and Fig 2A for a selection of snapshots. The statistical analysis of topography shape for the dynamics of five selected HECT molecule data sets is shown in Fig 2B. As we find, the automated shape analysis allows a classification of individual molecule dynamics: dwelling in round and oval shape states, population of intermediate shape topographies, and populations centered around either round or oval shape states.

**Fig. 2:**
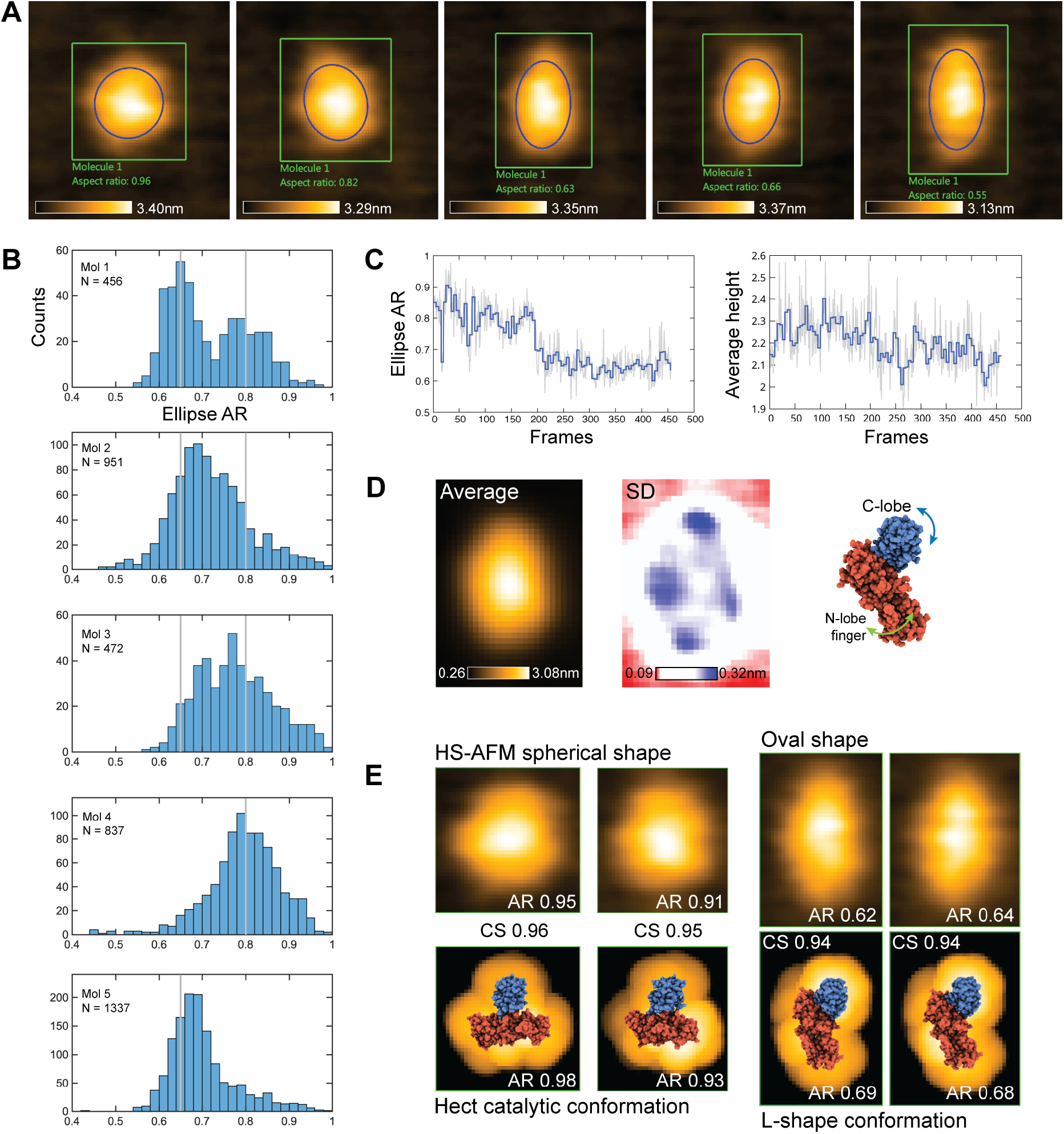
Automated molecule detection and statistical analysis of topography shape. Applications to HS-AFM imaging of the HECT domain protein. A: Exemplary snapshots demonstrating automated detection of dynamic topographies and quantitative shape analysis by ellipse fitting. B: Statistical analysis of topography shape shown as histograms of the ellipse fit aspect ratio (AR) for the recorded dynamics of 5 selected Hect individual molecules (number of frames N). Grey bars indicate AR values representative of round or oval shape, chosen from the analysis of molecule 1. C-E: Detailed analysis for dynamics of Mol 1. C: Time recording of the ellipse fit AR and the average height. D: Average AFM topography of molecule shapes aligned across all image frames (left); corresponding heat map with colors indicating standard deviation of heights across the pixel grid (middle); atomistic structure of the HECT domain L-shape conformation (PDB 1d5f) in a putative consistent alignment (right). E: Selected HS-AFM snapshots of spherical and oval topography shapes (top), and molecular orientations superimposed with their simulated AFM topographies (bottom) obtained from rigid-body fitting of HECT catalytic conformation (PDB 3jvz) and L-shape state. Corresponding ellipse AR values and image similarity scores (cosine similarity, CS) are provided.

For the exemplary case of a HECT molecule with a bi-modal distribution of the ellipse aspect ratio (molecule 1), the aspect ratio time trace shows that the dynamics of this molecule is characterized by a step-like transition from round shape to oval shape topographies (Fig 2C). During the time course the average topography height does not significantly change, indicating that the observed dynamical changes proceed parallel to the AFM substrate surface.

Our previous analysis of HECT domain HS-AFM imaging employing molecular modelling [15] and flexible fitting [11] of available atomistic structures showed that the flexibility of a single compact domain (C-lobe) and that of an extended finger domain is underlying the conformational transition between symmetric catalytic HECT structure (round AFM topographies) and L-shape structure (oval AFM topographies).

The new data evaluated here is consistent with this picture, as we demonstrate using information from the heat map of topography dynamics obtained from automation (Fig 2D) and rigid body fitting (Fig 2E) within the BioAFMviewer.

As a further application of molecule detection, we considered previously obtained data sets for HS-AFM imaging of the Atg1 protein which consists of two globular domains connected by an intrinsically disordered flexible linker. Before, the observed dynamics of Atg1 was analyzed by manual detection of the two domain linker junctions in all HS-AFM images to estimate the end-to-end distance of the flexible linker [16]. Here, we demonstrate that similar analysis results are obtained by a completely automatized workflow collecting statistics for several thousands of images (Fig. 3). The two global domains are identified by molecule detection with their distance being recorded (see Methods). Limitations of this automation include cases in which pairing of the two domains fails due to the presence of another molecule detected within the observation area, one of the molecules cannot be detected, or the fusion of both molecules (see Fig. 3B).

**Fig. 3:**
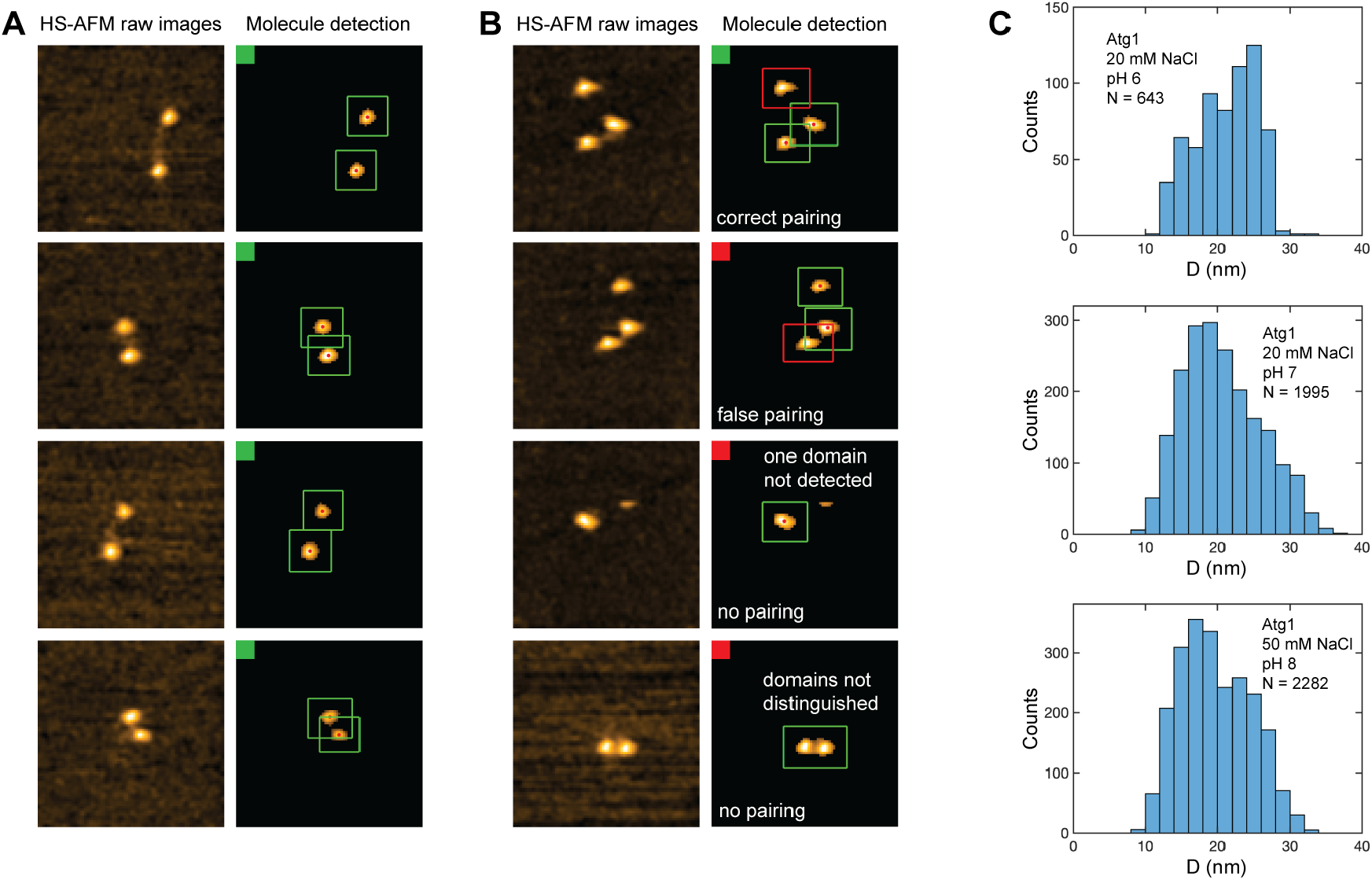
Automated molecule detection and statistical analysis of conformational dynamics. Application to HS-AFM imaging data of Atg1 protein dynamics. A: Snapshots representing the majority of reliable domain assignments to record end-to-end distance statistics. Side-by-side view of HS-AFM images and corresponding denoised images after molecule detection. Green boxes visualize the molecule region of interest; red dots indicate the molecule center of mass B: Exemplary cases of ambiguous and failed domain assignment. Red boxes show the presence of a third detected molecule. C: Histograms of the end-to-end distance of the Atg1 protein for three different buffer conditions obtained from fully automated analysis.

### 3. Dynamic molecule tracking

As a next step towards automation of AFM image analysis, we developed and implemented a method for dynamic tracking of detected molecules. Tracking is important to examine multiple molecules which are simultaneously observed while maintaining correct labeling of their identity to allow for high-throughput statistical analysis of imaging data. Details of the tracking algorithm can be found in the Methods section.

We considered two different HS-AFM data sets visualizing the dynamics of multiple HECT domains. In the software, motions of individual molecules are visualized as colored trajectories throughout the HS-AFM movie. The measurements and statistics are graphically displayed in the results panel for each molecule and can be combined into cumulative analysis.

Tracking of six individual HECT domain molecules over a time span of ∼32 sec (130 imaging frames) is demonstrated in Supplementary Movie V2. Fig. 4A shows selected example snapshots. At some moment, a seventh molecule enters the scanning area which is recognized and tracked by our algorithm (Fig. 4A,d). The algorithm is capable of tracking molecules even along irregular trajectories, allowing individual molecules to be followed despite substantial displacement from their initial position (molecules 2 and 4 in Fig. 4A; Suppl. Movie V2).

**Fig. 4:**
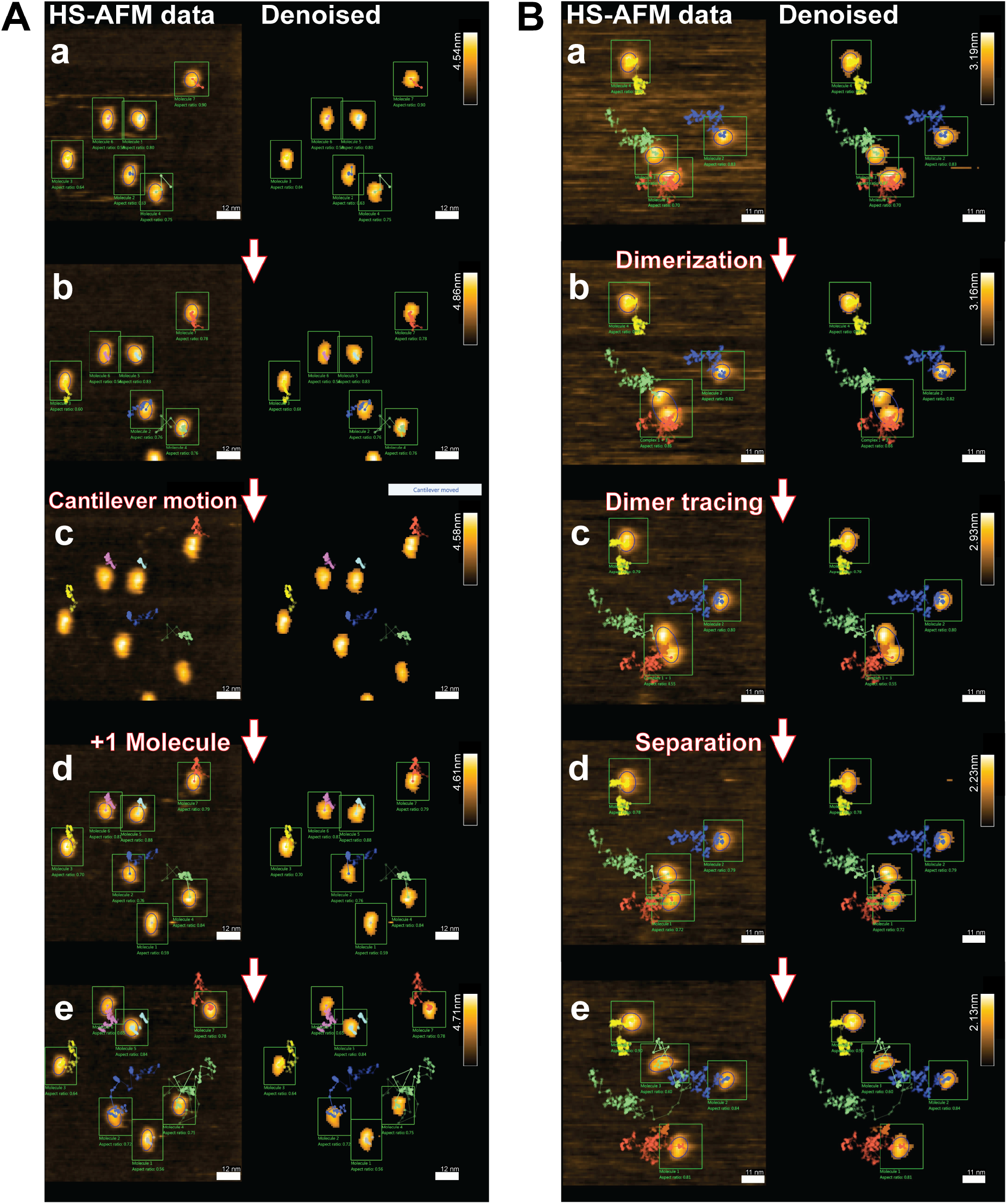
Molecule tracking. Application of molecule tracking to two separate HS-AFM imaging data sets of HECT domain dynamics. HS-AFM images and corresponding denoised images are shown side-by-side. Detected molecules are numbered and outlined by green boxes (ROI). Aspect ratio (AR) values of ellipse fitting are given. Dynamic tracking of individual molecule motion is visualized as traces of molecule topography centers in line-dot style using different colors. A: Events of cantilever motion and addition of a HECT molecule to the AFM canvas are indicated to highlight versatility and robustness of the tracking algorithm. B: Events of molecule dimerization, dynamic tracking of the dimer, and its separation are highlighted.

Tracking for the second examined example of four individual HECT domain molecules over a time span of ∼52 sec (471 imaging frames) is demonstrated in Supplementary Movie V3 with some exemplary snapshots shown in Fig. 4B.

This example includes a highly mobile molecule approaching another molecule followed by subsequent dimerization (Fig. 4B,b). This dimerization process is detected, and a new trajectory is created and tracked for the newly formed dimer (Fig. 4B,c). After some time, the two molecules dissociate and are reassigned to their original individual trajectories (Fig. 4B (d,e)). Both examples demonstrate the robustness of our tracking algorithm to handle complex situations including changes in molecule number and following relative motions between molecules taking place over several tens of nm.

To demonstrate versatility of molecule detection combined with tracking we next considered HS-AFM observations of a lipid bilayer formed on a mica substrate which showed large areas which correspond to unfilled holes (Fig. 5). To perform image segmentation separating membrane and holes we have implemented often used inversion of topography heights (see Methods) to generate inverted AFM images in which holes appear as elevated globular objects which can be detected and tracked as molecules by the same described approach.

**Fig. 5:**
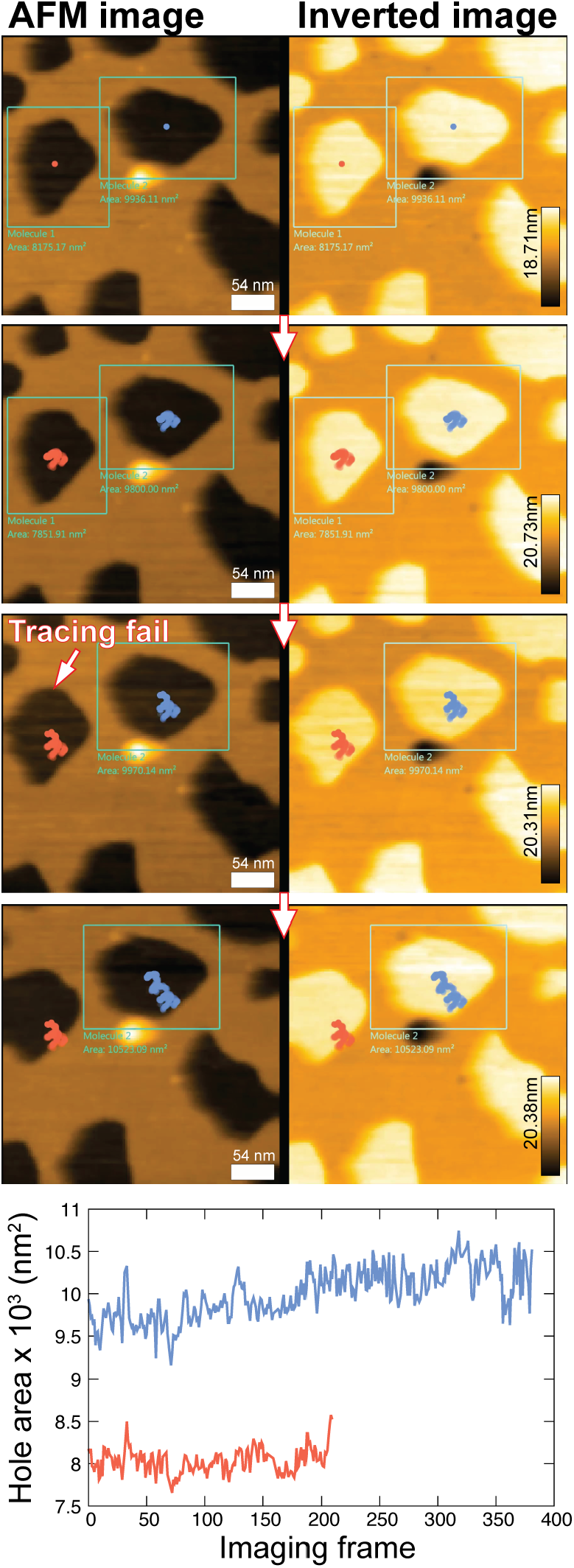
HS-AFM image segmentation and automated tracking. Automated image segmentation of HS-AFM data for a lipid membrane covered mica surface and dynamic tracking of void areas by the application of imaging height inversion. Original AFM images and corresponding inverted images are shown side-by-side. The two regions detected as void regions are outlined as boxes (ROI), values of respective topography area are provided, and center traces are visualized in red/blue colors. The case where detection and tracing fails is indicated. Time traces for the topography area of the two void regions are shown at the bottom.

The application of this method is demonstrated in Supplementary Movie V4. Selected snapshots of original and inverted HS-AFM images are shown in Fig. 5, including quantitative analysis by monitoring time traces of hole area (Fig. 5). While in the presented case the temporal increase of area is likely attributed to degradation of lipids at hole boundaries induced by scanning tip force, applications of tracking and quantitative analysis can be further employed to understand phenomena of active degradation by enzymes and antibiotics [17,18] or temperature induced restructuring [19] of model membranes observed by AFM.

Within the context of tracking, we also provide a demonstration of post-experimental analysis to remove parachuting effects from HS-AFM imaging which we implemented into our software based on a previously developed method [20]. The application to a HS-AFM movie of four HECT domain molecules is demonstrated in Supplementary Movie V5 and selected snapshots are shown in Suppl. Fig. S1. The effect of image correction was quantified by tracking the area of detected molecules.

## 4. Conclusion

With the BioAFMviewer AFM Analysis Arena (A^3^) we provide a unique software platform addressing the demand for automated high-throughput analysis of HS-AFM imaging data towards transforming measurements from observations into an analytical tool to improve the quantitative understanding of nanoscale biological processes.

The user-friendly interactive interface providing a direct workflow from raw experimental data to statistical analysis opens the opportunity for immediate practical applications by the AFM community.

## 5. Software availability

The BioAFMviewer software is available as a free download from the project website www.bioafmviewer.com. From version 6.0 onwards, the software is divided into two interfaces. The first, called Nano AFM Integrative Modeling and Simulation (NanoAIMS), is the platform for integrative modeling and simulation and includes the functionalities developed in previous versions of the software. These include an interactive interface for live simulation AFM of available structural data [10], rigid-body fitting to AFM images [21], molecule placement prediction based on electrostatic interactions [12] and the implementation of Normal Mode Flexible Fitting AFM (NMFF-AFM) [11]. The second interface, AFM Analysis Arena (A3), includes all the developments introduced in the present work.

## Acknowledgements

We are grateful to Noriyuki Kodera for providing the HS-AFM data of the Atg1 protein and of the lipid bilayer formed on mica. We thank Shintaroh Kubo and Shoji Takada for sharing the source code used to remove parachuting from HS-AFM imaging data. We also thank Kazusa Takeda for providing the HS-AFM data of HECT domain used to demonstrate the application of parachuting removal. This work was supported by the Ministry of Education, Culture, Sports, Science and Technology (MEXT), Japan, through the World Premier International Research Center (WPI) Initiative. This work was also supported by WPI-NanoLSI Transdisciplinary Research Promotion Grants, Kanazawa University.

## Supplementary Material

**Supporting Table T1:** Shape descriptors. Overview of descriptors implemented in A3 to quantify shape of detected molecules from AFM imaging.

| Shape descriptors | Measurement | Dimension |
| --- | --- | --- |
| Free form shape |  |  |
| Circle fit | Radius (R) | nm |
| Ellipse fit | Major ($R_M$) & minor ($R_m$) axes lengths | nm |
| Area (A) | $\sum_{\text{pix} \in \text{mol}} (\text{scan step})^2$ | $\text{nm}^2$ |
| Volume (V) | $\sum_{\text{pix} \in \text{mol}} (h - \bar{h}_{\text{BKG}}) \cdot (\text{scan step})^2$ | $\text{nm}^3$ |
| Average height | $\sum_{\text{pix} \in \text{mol}} h / \sum_{\text{pix} \in \text{mol}} 1$ | nm |
| Maximum height | $\max(\{h\}_{\text{pix} \in \text{mol}})$ | nm |
| Perimeter (P) | $\sum_{\text{pix} \in \text{mol edge}} \text{scan step}$ | nm |
| Aspect ratio | $R_m/R_M$ | 1 |
| Circularity | $4\pi A/P^2$ | 1 |
| Roundness | $4A/(\pi R_M^2)$ | 1 |
| Solidity | $A/(\text{convex hull area})$ | 1 |

**Supporting Table T2:** Data correlations. Table of Pearson correlation coefficients between available shape descriptors from the data set analysis of 20 different HECT molecules.

|  | Area | Volume | Avg. height | Max. height | Perimeter | Aspect ratio | Circularity | Roundness | Solidity | Radius |
| --- | --- | --- | --- | --- | --- | --- | --- | --- | --- | --- |
| Area | 1.00 | 0.79 | 0.40 | 0.40 | 0.85 | -0.08 | -0.19 | -0.09 | -0.08 | 0.77 |
| Volume |  | 1.00 | 0.83 | 0.82 | 0.55 | -0.08 | 0.05 | -0.04 | 0.15 | 0.56 |
| Avg. height |  |  | 1.00 | 0.95 | 0.20 | 0.00 | 0.17 | 0.04 | 0.22 | 0.24 |
| Max. height |  |  |  | 1.00 | 0.24 | -0.01 | 0.10 | 0.01 | 0.15 | 0.27 |
| Perimeter |  |  |  |  | 1.00 | -0.23 | -0.65 | -0.32 | -0.55 | 0.86 |
| Aspect ratio |  |  |  |  |  | 1.00 | 0.41 | 0.98 | 0.13 | -0.61 |
| Circularity |  |  |  |  |  |  | 1.00 | 0.56 | 0.90 | -0.56 |
| Roundness |  |  |  |  |  |  |  | 1.00 | 0.30 | -0.65 |
| Solidity |  |  |  |  |  |  |  |  | 1.00 | -0.33 |
| Radius |  |  |  |  |  |  |  |  |  | 1.00 |

**Supporting Fig. S1:**
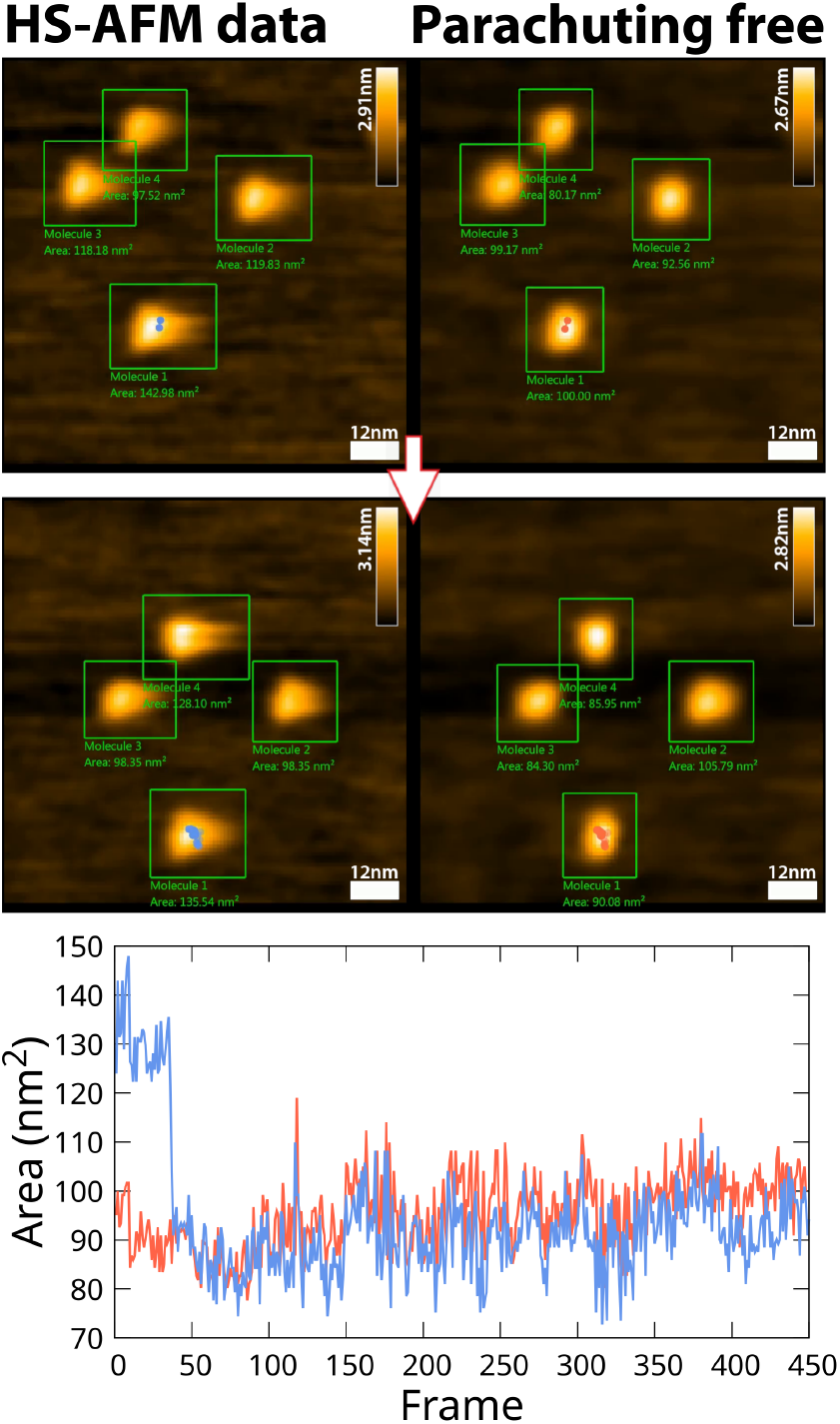
Parachuting removal. Application of the implemented parachuting removal method to HS-AFM imaging data of the HECT domain. Two exemplary snapshots from Movie V5 are shown. Values of the topography area are displayed for the 4 individual molecules. To quantitively demonstrate the effect, the topography area of detected molecule 1 recorded over time is displayed in blue (original data) and red color (parachuting free), respectively.

## Supplementary Material Videos

**Movie V1:** Automated molecule detection and tracing. Application of fully automated molecule detection, tracing, and dynamic shape quantification to HS-AFM imaging of single HECT domain. Shown is a side-by-side view of the raw HS-AFM data, the same data with application of a Gaussian filter, and the denoised version after Otsu thresholding. Ellipse fitting results are outlined in blue color with aspect ratio value being displayed. Traces of the molecule topography centers are visualized in line-dot style in red color.

**Movie V2:** Multiple molecules tracking ver 1. Application of molecule tracking to HS-AFM imaging data capturing topography dynamics of multiple HECT domain molecules. Shown is a side-by-side view of the raw HS-AFM data and the denoised version after Otsu thresholding. Detected molecules are numbered and outlined by green boxes (ROI). Aspect ratio (AR) values of ellipse fitting are given. Dynamic tracking of individual molecule motion is visualized as traces of molecule topography centers in line-dot style using different colors. Events of cantilever motion and addition of a HECT molecule to the AFM canvas highlight the versatility and robustness of the tracking algorithm.

**Movie V3:** Multiple molecules tracking ver 2. Application of molecule tracking to HS-AFM imaging data capturing topography dynamics of multiple HECT domain molecules. Shown is a side-by-side view of the raw HS-AFM data and the denoised version after Otsu thresholding. Detected molecules are numbered and outlined by green boxes (ROI). Aspect ratio (AR) values of ellipse fitting are given. Dynamic tracking of individual molecule motion is visualized as traces of molecule topography centers in line-dot style using different colors. Events of molecule dimerization, dynamic tracking of the dimer, and its separation are highlighted. Long-time tracking of individual molecules which undergo significant relative motions over the range of several tens of nm is demonstrated.

**Movie V4:** HS-AFM image segmentation and automated tracking. Automated image segmentation of HS-AFM data for a lipid membrane covered mica surface and dynamic tracking of void areas by the application of imaging height inversion. The HS-AFM original data movie and the version with applied height inversion are shown side-by-side. The two regions detected as void regions are outlined as boxes (ROI), values of respective topography area are provided, and center traces are visualized in red/blue colors.

**Movie V5:** Parachuting removal. Application of the implemented parachuting removal method to HS-AFM imaging data of 4 individual HECT domain molecules. The HS-AFM raw data movie and the version with applied parachuting removal are shown side-by-side. Values of the topography area are displayed for the 4 individual molecules.

